# Properties of genomic relationships for estimating current genetic variances within and genetic correlations between populations

**DOI:** 10.1101/124115

**Authors:** Yvonne C.J. Wientjes, Piter Bijma, Jérémie Vandenplas, Mario P.L. Calus

**Author notes:** Author information: Yvonne Wientjes, Wageningen University & Research, Animal Breeding and Genomics, P.O. box 338, 6700 AH Wageningen, the Netherlands.

## Abstract

Different methods are available to calculate multi-population genomic relationship matrices. Since those matrices differ in base population, it is anticipated that the method used to calculate the genomic relationship matrix affect the estimate of genetic variances, covariances and correlations. The aim of this paper is to define a multi-population genomic relationship matrix to estimate current genetic variances within and genetic correlations between populations. The genomic relationship matrix containing two populations consists of four blocks, one block for population 1, one block for population 2, and two blocks for relationships between the populations. It is known, based on literature, that current genetic variances are estimated when the current population is used as base population of the relationship matrix. In this paper, we theoretically derived the properties of the genomic relationship matrix to estimate genetic correlations and validated it using simulations. When the scaling factors of the genomic relationship matrix fulfill the property 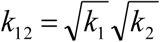, the genetic correlation is estimated even though estimated variance components are not necessarily related to the current population. When this property is not met, the correlation based on estimated variance components should be multiplied by 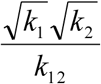 to rescale the genetic correlation. In this study we present a genomic relationship matrix which directly results in current genetic variances as well as genetic correlations between populations.

## INTRODUCTION

When estimating additive genetic values of individuals, the relationships between individuals are used to describe the covariance between additive genetic values for a specific trait. Those covariances between individuals are best represented by the relationships at causal loci. Since causal loci are generally unknown, different approaches have been developed to estimate relationships from genomic marker data (e.g., VanRaden 2008; Powell *et al*. 2010; Yang *et al*. 2010). As long as causal loci and genomic markers have the same properties, such as allele frequency distribution, relationships at the markers are observed to be an unbiased estimate of relationships at the causal loci (Yang *et al*. 2010; Yang *et al*. 2015).

Relationships are expressed relative to a base population, consisting of unrelated individuals that have average self-relationships of one, for which the additive genetic variance is estimated. The base population of a genomic relationship matrix depends on the method used to calculate the relationship matrix, therefore estimated variances differ across methods (Speed and Balding 2015; Legarra 2016). By using the current allele frequencies to calculate the genomic relationship matrix, the current population is the base population for which additive genetic variances are estimated (Hayes *et al*. 2009).

Genomic relationships can also be calculated between distantly related individuals, for example between individuals from different populations. Those relationships can be used to estimate genetic correlations between populations using a multi-trait model (Karoui *et al*. 2012), where the same trait in each population is modelled as a different trait. Due to differences in environments and allele frequencies, in combination with non-additive effects, the allele substitution effects of causal loci can differ between populations (e.g., Fisher 1918; Fisher 1930; Falconer 1952). Moreover, some causal loci might only segregate in one of the populations. Therefore, the genetic correlation between populations can differ from 1.

The genetic correlation between populations is an important parameter, since it is used to understand the genetic architecture and evolution of complex traits, such as disease traits in humans (De Candia *et al*. 2013; Brown *et al*. 2016). Moreover, the genetic correlation determines whether information can be shared across populations as done in multi-population genomic prediction (Wientjes *et al*. 2015; Wientjes *et al*. 2016), which is of importance for animals (e.g., Karoui *et al*. 2012; Olson *et al*. 2012), plants (e.g., Lehermeier *et al*. 2015) and humans (e.g., De Candia *et al*. 2013).

Different methods are available to calculate multi-population genomic relationship matrices (Harris and Johnson 2010; Erbe *et al*. 2012; Chen *et al*. 2013; Makgahlela *et al*. (2013). The two most important differences between the methods are: 1) the assumed relation between effect size and allele frequency of markers; namely assuming effect size and allele frequency are independent (e.g., method 1 of VanRaden (2008)) or assuming that markers with a lower allele frequency have a larger effect (e.g., method 2 of VanRaden (2008) and Yang (2010)), and 2) the allele frequency that is used; namely allele frequencies specific to each population, the average allele frequency across the populations, or the estimated allele frequency when the populations separated. Since relationships between individuals differ across those methods, it is anticipated that the method used to calculate the genomic relationship matrix affects the estimate of the genetic correlation.

Therefore, the aim of this paper is to define a multi-population genomic relationship matrix to estimate current genetic variances within and genetic correlations between populations. We theoretically derive a relationship matrix with this property and validate it with simulations. To rule out the effect of differences in linkage disequilibrium between markers and causal loci, we will focus in the entire paper on a situation where causal loci are used to calculate the relationships.

## MATERIALS AND METHODS

### Theory

The additive genetic correlation, *r_g_*, is the correlation between additive genetic values (*A*) for two traits of the same individual (Bohren *et al*. 1966; Falconer and Mackay 1996). In an additive model and under the assumptions that the correlation originates from pleiotropy, genetic values are independent between loci, and allele substitution effects are independent from allele frequency, *r_g_* is equal to the average correlation between allele substitution effects of the two traits, denoted as trait 1 and 2, at causal loci. This equality can be shown for individual *i* by considering both genotypes (*z*) and allele substitution effects (α) at all *n_c_* causal loci as random:

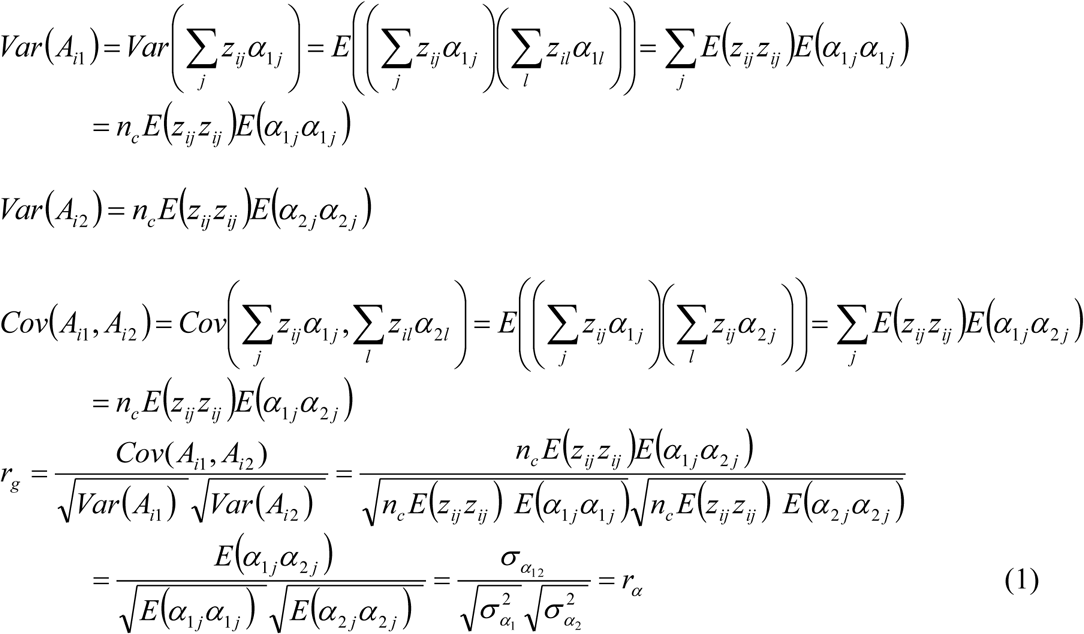
where *j* and *l* denote the different causal loci. Genotypes are represented by allele counts coded as 0, 1 and 2 that are centered by subtracting *2p*, where *p* is the allele frequency for the counted allele.

Similar to genetic correlations between traits in one population, the genetic correlation (*r_g_*) between populations can be estimated in a multi-trait model using a relationship matrix and REML by modelling the phenotypes of two populations as different traits (Karoui *et al*. 2012). This approach is also known as multi-trait GREML. In the following, we will refer to trait 1 as the trait expressed in population 1 and to trait 2 as the trait expressed in population 2. When considering performance in different populations as different traits, individuals have a phenotype for only one trait. Therefore, the (co)variance structure of the additive genetic values can be written as (Visscher *et al*. 2014):

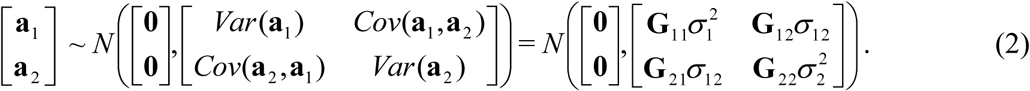
where **a**_1_ is the vector with additive genetic values for individuals from population 1 for trait 1, **a**_2_ is the analogous vector for individuals from population 2 for trait 2, 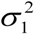 and 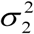 are genetic variances for the two traits, *σ_12_* is the genetic covariance between the traits, **G**_11_ is a matrix with genomic relationships within population 1, **G**22 is a matrix with genomic relationships within population 2, and **G**_12_ and 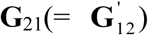 are matrices with genomic relationships between population 1 and 2.

To derive the definition of the genomic relationships in Equation 2, we derive the variances and covariance of the additive genetic values for the two traits. Naturally, this will result in an equation to calculate the genomic relationship matrix (**G**) across populations to estimate (co)variances in the current populations.

When both populations are in Hardy-Weinberg equilibrium, allele substitution effects are independent from allele frequency, and effects of causal loci are independent from each other, the genetic variance for trait 1 can be written as 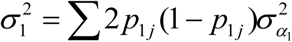, where *p_1j_* is the allele frequency at locus *j* in population 1 (Falconer and Mackay 1996). Hence, the variance of **a**1 is:

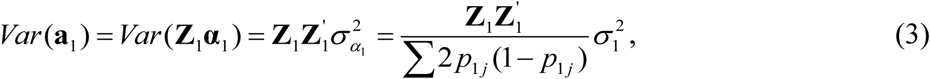
where **Z**_1_ is a *n_1_* x *n_c_* matrix of centered genotypes for all individuals from population 1 (*n_1_*) for all causal loci, and **α**_1_ is a vector of length *n_c_* with allele substitution effects at causal loci for trait 1.

Similarly,

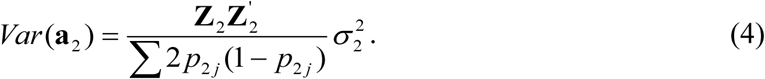

The genetic covariance between the two traits is:

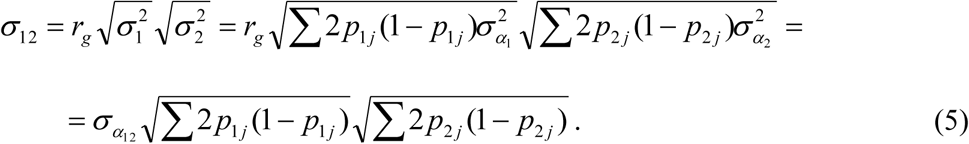
Therefore, the covariance between genetic values of population 1 and 2 is:

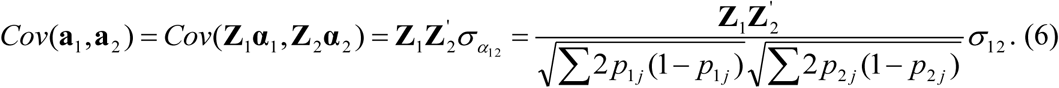
From Equation 3, 4 and 6, it follows that the genomic relationship matrix (**G**) is:

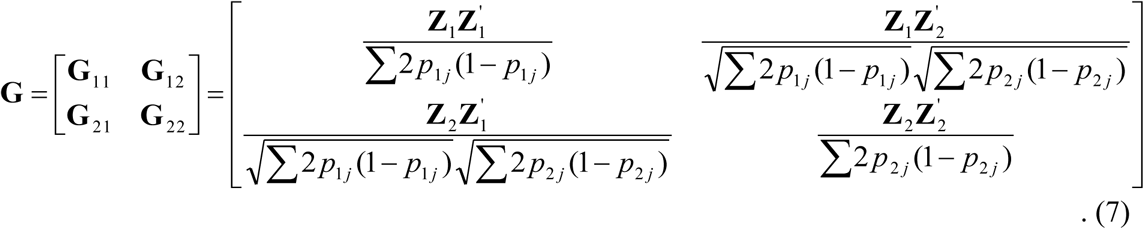
When allele frequencies from the current population are used, **G** from Equation 7 estimates current genetic (co)variances. Lourenco *et al*. (2016) presented a comparable **G** matrix for combining purebred and crossbred animals. Note that the covariance of the genotypes between the populations, 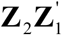, is divided by the standard deviations of the genotypes in each population, 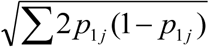 and 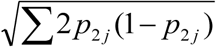. Therefore, the relationships in this **G** matrix are defined as correlations between the individuals.

By interpreting 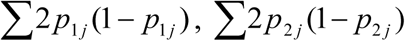 and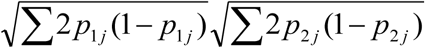 as scaling factors (i.e. *k_1_, k_2_*, and *k_12_*) of **G**, the variance-covariance matrix in Equation 2 becomes:

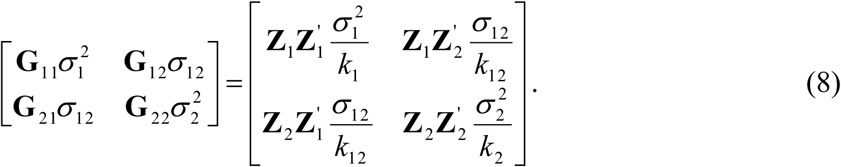
Equation 8 shows that the scaling factors of **G** and the variance components are completely confounded. Therefore, other scaling factors of **G** can be used to estimate the genetic correlation as:

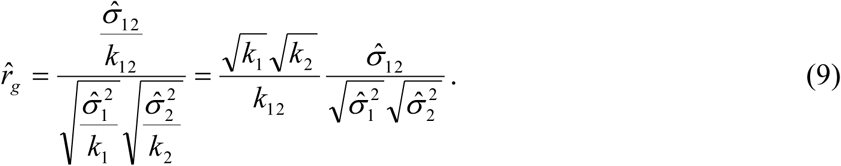
Equation 9 shows that the genetic correlation is directly estimated from the variance components when the scaling factors of **G** fulfil the property 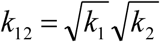. When 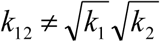, the correlation based on variance components should be multiplied by 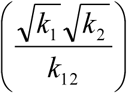 to correct the estimated genetic correlation. By changing the scaling factors, the genetic variances change as well. When genetic variances of the current population are of interest, the within-population blocks in **G** should be scaled as in Equation 7 (Legarra 2016).

Equation 8 and 9 show that the genetic correlation is estimated when the scaling factors in **G** are the same for all blocks. When all scaling factors are equal to 1, so effectively no scaling factor is used, the (co)variances represent the (co)variances of the causal effects i.e., 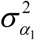, 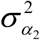, and *σ_α_12__*. A disadvantage of this scaling is that elements of **G** can become very large, which can result in very small variance components that may be flagged as too small in statistical software. This might be prevented by either scaling up the phenotypic variance by multiplying all phenotypes by a constant, or by scaling down the elements in **G** by dividing all elements by the same constant. Both scaling approaches have no influence on the genetic correlation, but do affect the genetic (co)variances.

### Simulations

Simulations were used to validate the results above. Two populations of 2500 individuals each with phenotypes for a trait influenced by the same 15 000 loci were simulated. Allele frequencies of the loci were sampled from a U-shape distribution, independently in both populations. Genotypes were allocated to individuals according to Hardy-Weinberg equilibrium, assuming that loci were segregating independently. Therefore, genetic correlations between populations were only affected by pleiotropy and not by linkage disequilibrium.

Allele substitution effects were sampled from a bi-variate normal distribution with means zero and variances 1, and a correlation of 0.5 between allele substitution effects in both populations. The allele substitution effects were multiplied with the corresponding genotypes to calculate additive genetic values for individuals, assuming additive gene action. Environmental effects were sampled from a normal distribution with variance 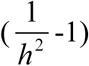 times the genetic variance, where the genetic variance was calculated across all individuals in both populations. The heritability was set to 0.9, to ensure that there was sufficient power in the data to estimate the (co)variances. Phenotypes were the sum of additive genetic and environmental effects, and were standardized to an average of 0 and a standard deviation of 100. Simulations were replicated 100 times.

Phenotypes were analyzed in a two-trait model, using four different **G** matrices; two **G** matrices derived above, and two commonly used **G** matrices for multiple populations (Chen *et al*. 2013; Makgahlela *et al*. 2013). In all four methods, genotypes at causal loci were used to calculate **G.** The methods differed in scaling factors as well as in centering of genotypes, being performed either within or across populations.

In the first three methods, the genotypes in **Z** were centered within population as *g_ijm_* –2*P_jm_*, where *g_ijm_* is the allele count of individual *i* from population *m* at locus *j* and *p_jm_* is the allele frequency at locus *j* in population *m*. The first method, **G**_New, scaled **G** following Equation 9:

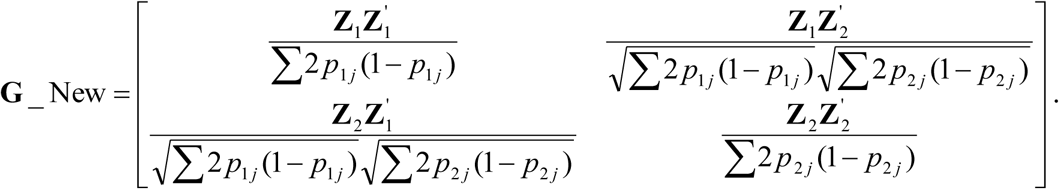
In the second method, **G**_1, scaling factors were equal to 1:

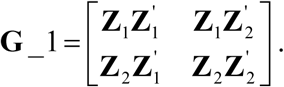
The third method, **G**_Chen, calculated **G** according to Chen et al. (2013):

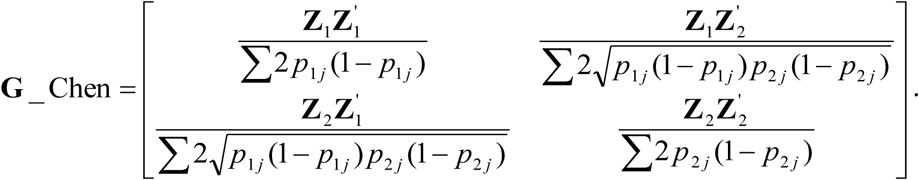

The fourth method, **G**_Across, used the average allele frequency across both populations instead of population-specific allele frequencies to center the genotypes (e.g., Makgahlela *et al*. 2013). Thus, the matrix of genotypes, denoted **Z\***, had elements 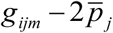, where 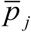 is the average allele frequency across both populations at locus *j*. The scaling factor was the same for all blocks:

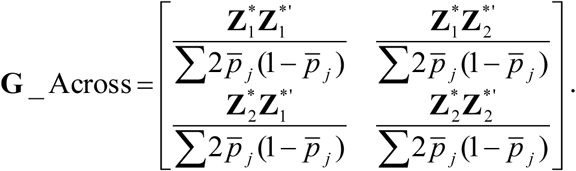

**G**_New, **G**_1 and **G**_Across fulfilled the property 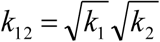 to directly estimate the genetic correlation. In **G**_Chen, 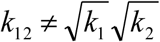 when allele frequencies in the two populations were different. Therefore, the genetic correlation estimated with **G**_Chen was multiplied by 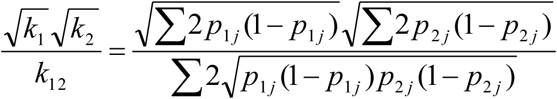 to correct the estimate. Moreover, the current populations were the base population for the within-population blocks of **G**_New and **G**_Chen, so those **G** matrices estimated the genetic variances within the current populations (Speed and Balding 2015; Legarra 2016). As explained before, the variances of **G**_1 represented the variances of the causal effects. For **G**_Across, the base population was not clearly defined, so the interpretation of the estimated genetic variances is unclear. See supporting information for the R-script and seeds used to simulate genotypes and phenotypes and to calculate the different **G** matrices.

## RESULTS

### Variance components

In Figure 1, the estimated genetic variance using **G**_New is plotted against the simulated genetic variance. This figure shows that the estimates varied only slightly around the simulated values. This shows that **G**_New unbiasedly estimated the genetic variance in the current populations.

**Figure 1.**
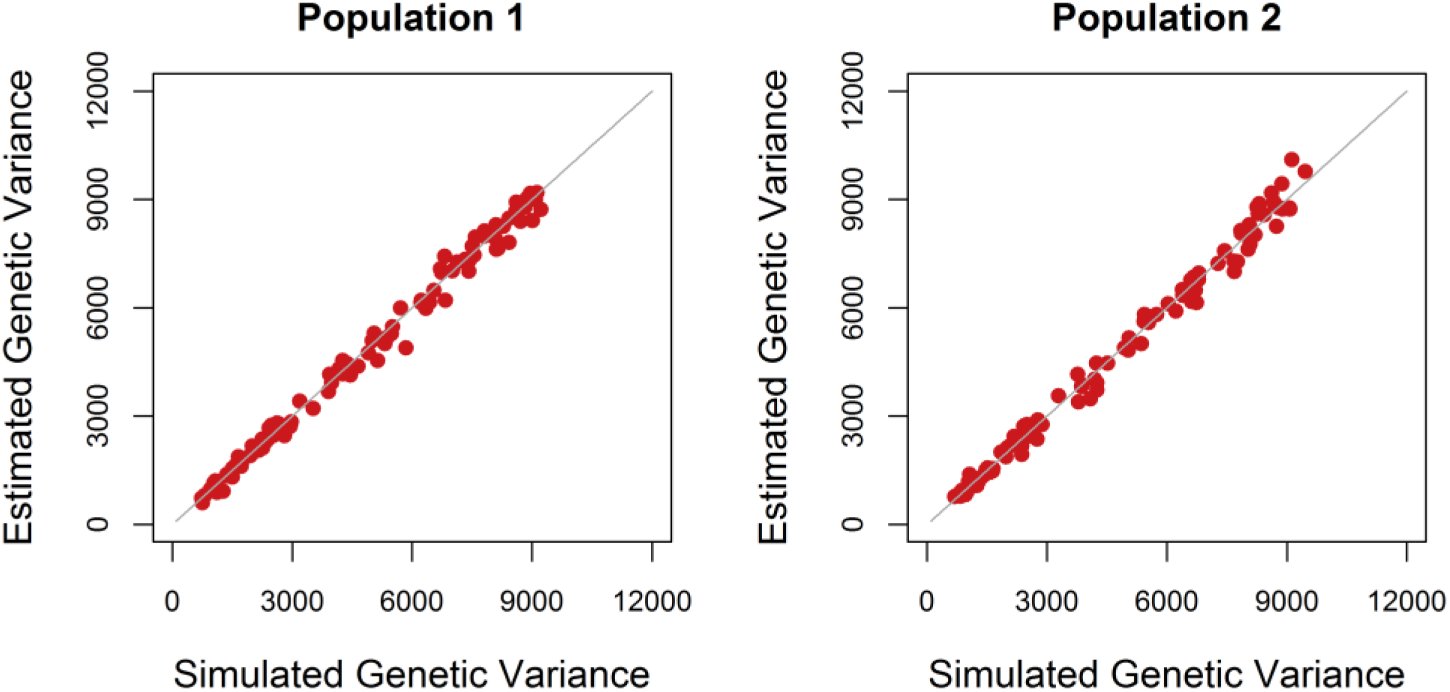
Estimated versus simulated genetic variance. The estimated genetic variance in both populations in each of the 100 replicates using the genomic relationship matrix derived in this study (**G**_New) versus the simulated genetic variance. The grey line represents the line y=x.

As expected, **G**_New and **G**_Chen estimated the same genetic variances (Figure 2 and 3). The variances of **G**_1 represented the variances of the causal effects. By multiplying those variances by ∑2*P_jm_*(1 – *P_jm_*) for population m, genetic variances identical to **G**_New and **G**_Chen were obtained. The genetic variance estimated with **G**_Across was approximately a factor 1.5 higher than the genetic variance estimated with **G**_New and **G_**Chen. Also the scaling factors *k_1_* and *k_2_* were approximately a factor 1.5 higher. Hence, when multiplying the variances estimated with **G**_Across by the ratio in scaling factors, estimates became identical to those with **G**_New and **G**_Chen. So, the difference in estimated variances between methods was completely explained by the difference in scaling factors, while centering genotypes within or across populations had no effect on estimated variances. Estimated residual variances were exactly the same for the four different **G** matrices.

**Figure 2.**
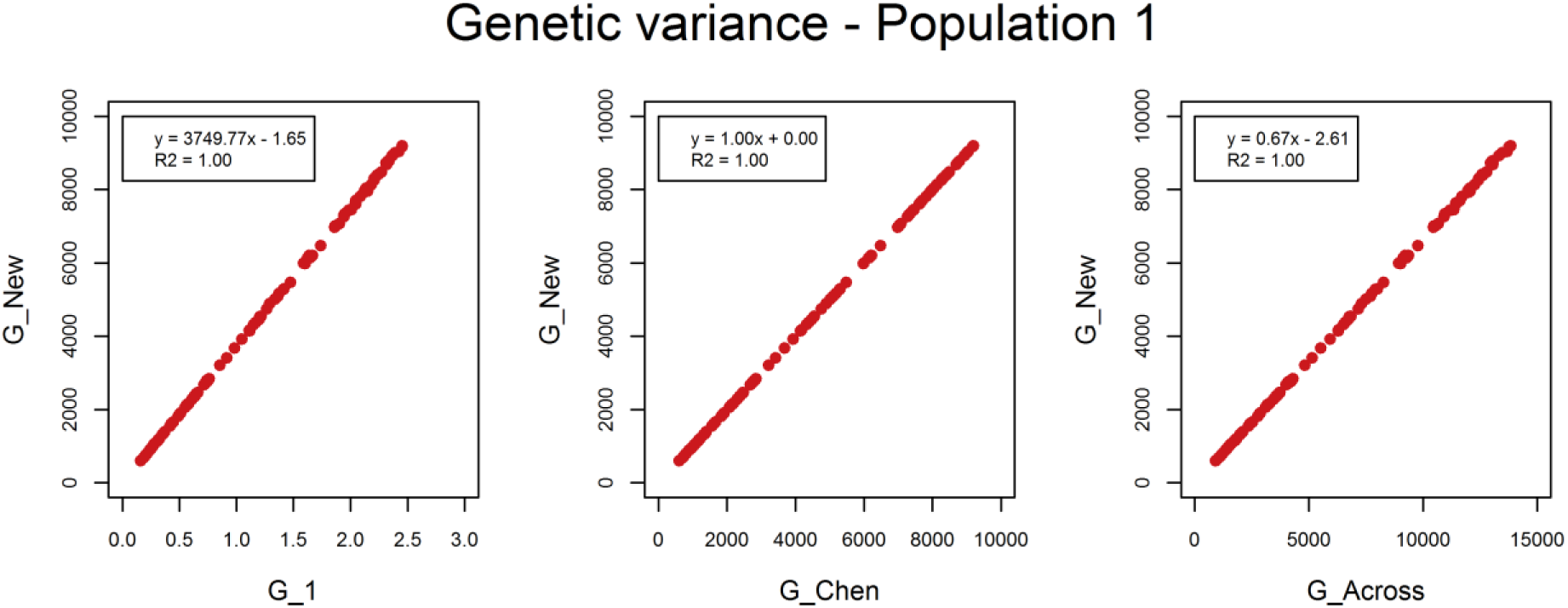
Estimated genetic variance in population 1. The estimated genetic variance in population 1 in each of the 100 replicates using the genomic relationship matrix derived in this study (**G**_New) versus the estimated genetic variance using population-specific allele frequencies and either a genomic relationship matrix without scaling factors (**G**_1) or based on the method of Chen *et al*. (2013; **G_**Chen), or using allele frequencies across populations (**G**_Across).

**Figure 3.**
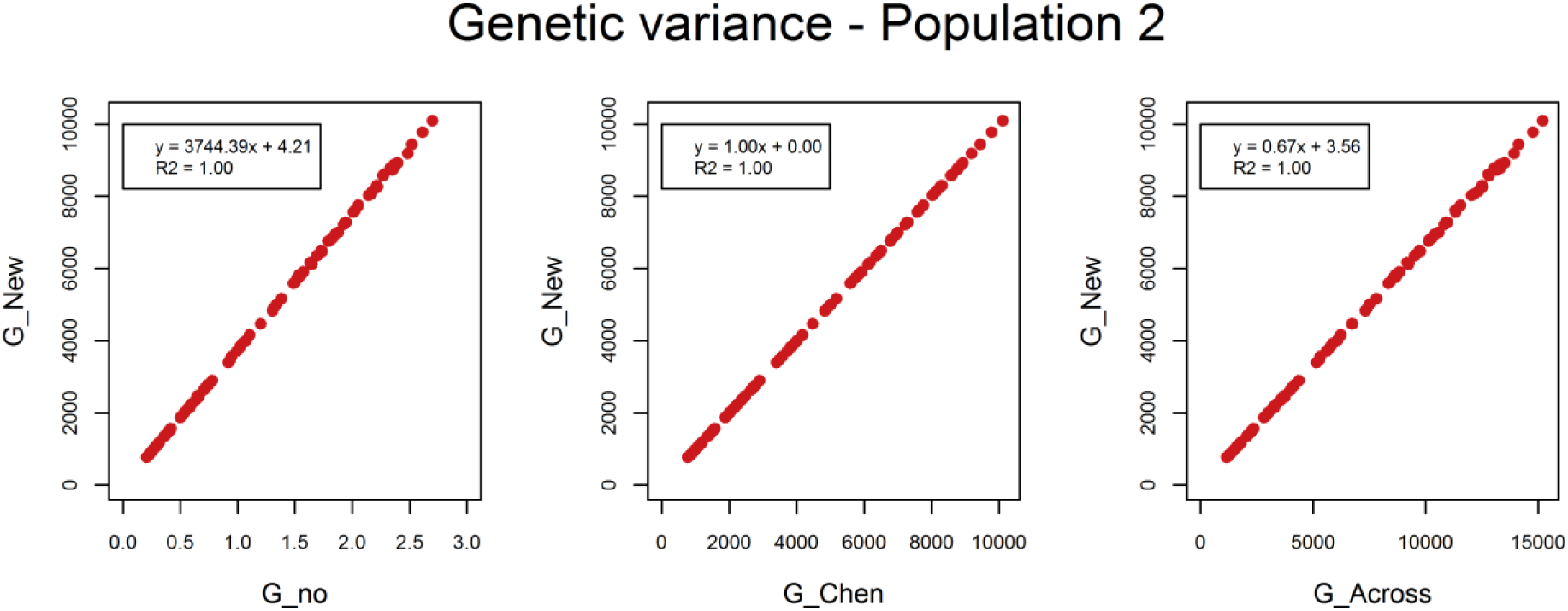
Estimated genetic variance in population 2. The estimated genetic variance in population 2 in each of the 100 replicates using the genomic relationship matrix derived in this study (**G**_New) versus the estimated genetic variance using population-specific allele frequencies and either a genomic relationship matrix without scaling factors (**G**_1) or based on the method of Chen *et al*. (2013; **G**_Chen), or using allele frequencies across populations (**G**_Across).

### Genetic correlation

Despite differences in (co)variance estimates, **G_**New, **G_**1, and **G**_Across yielded the same estimated genetic correlation (Figure 4) which was an unbiased estimate of the simulated genetic correlation (Figure 5). This is because differences in genetic covariances among models were compensated by corresponding differences in genetic variances. The genetic correlation estimated using **G**_Chen was ∼20% lower. When multiplying this estimate by 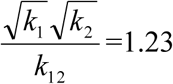, the genetic correlation became identical to the other three methods.

**Figure 4.**
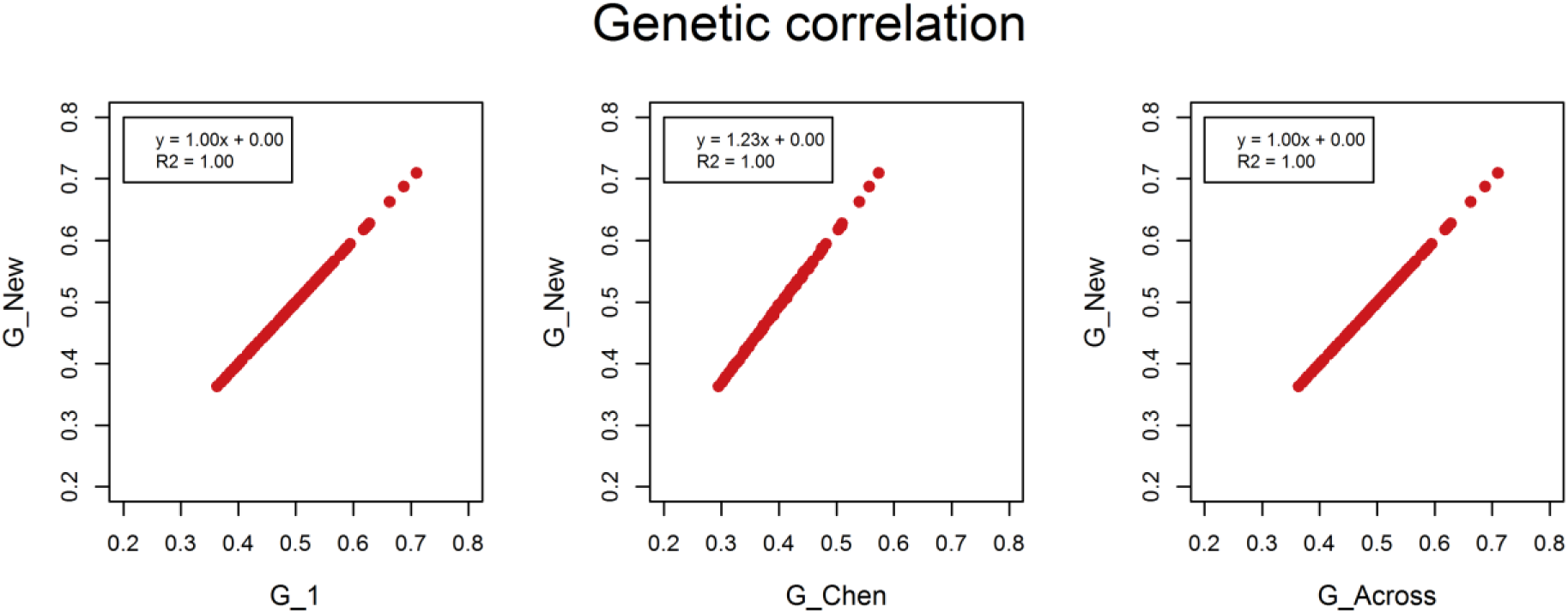
Estimated genetic correlation between population 1 and 2. The estimated genetic correlation between population 1 and 2 in each of the 100 replicates using the genomic relationship derived in this study (**G**_New) versus the estimated genetic correlation using population-specific allele frequencies and either a genomic relationship matrix without scaling factors (**G**_1), based on the method of Chen *et al*. (2013; **G**_Chen), or using allele frequencies across populations (**G**_Across).

**Figure 5.**
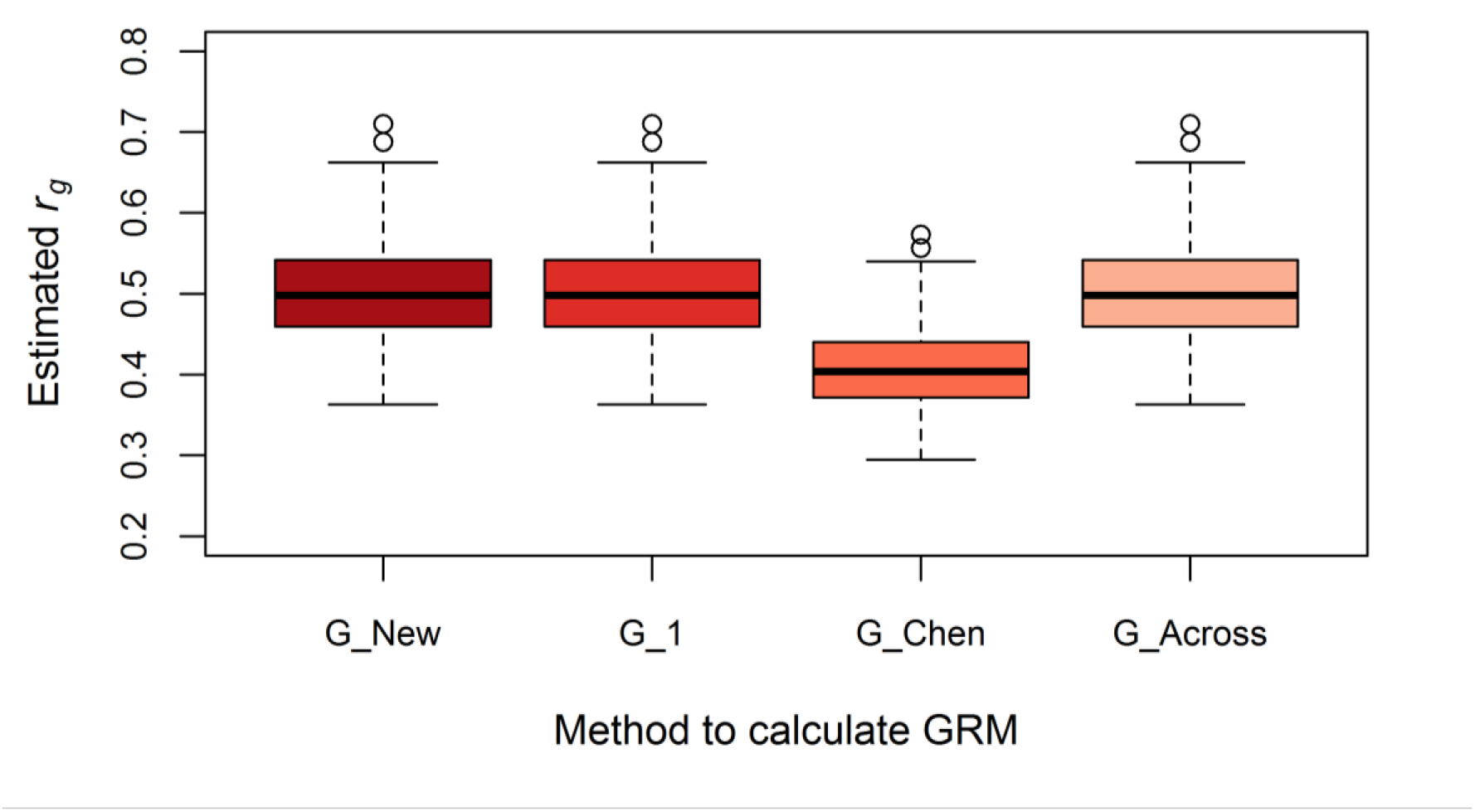
Boxplot of the estimated genetic correlation using different methods to calculate the genomic relationship matrix. The estimated genetic correlation between population 1 and 2 in each of the 100 replicates using the genomic relationship matrix derived in this study (**G**_New), using population-specific allele frequencies and either a genomic relationship matrix without scaling factors (**G**_1), or based on the method of Chen *et al*. (2013; **G**_Chen), or using allele frequencies across populations (**G**_Across). The simulated genetic correlation was 0.5.

## DISCUSSION

The aim of this paper was to define a multi-population genomic relationship matrix to estimate current genetic variances within and genetic correlations between populations. We derived a genomic relationship matrix, **G**_New, that yields unbiased estimates of current genetic variances, covariances and correlations. Moreover, we showed the required property for other genomic relationship matrices to estimate the genetic correlation between populations, even though estimated variance components are not necessarily related to the current populations.

### Methods to calculate the genomic relationship matrix

From the four methods used in this paper to calculate **G**, **G**_New was the only matrix correctly estimating both current genetic variances as well as genetic correlations. **G**_Chen also estimated current genetic variances, but the estimated genetic correlation had to be multiplied by 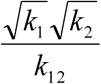. **G**_1 estimated the correct genetic correlation, but estimated the variance of causal effects instead of the genetic variance. Although the base population in **G**_Across was not well defined, genetic correlations were correctly estimated but there was no clear interpretation of the estimated genetic variances. Results also showed that genetic variances were not affected by centering the allele count, as shown before by Strandén and Christensen (2011).

Table 1 gives an overview of the most frequently used methods to calculate **G** across multiple populations, with scaling factors and correction factors for the estimated genetic correlation. **G**_New, **G**_1, **G**_Across, and the method described by Erbe *et al*. (2012) directly estimate the correct genetic correlation. The **G_**Chen method does not directly estimate the genetic correlation, but the estimate can be corrected using the scaling factors. Those five methods all assume that allele substitution effects are independent of allele frequency, similar to method 1 of VanRaden (2008). This is in contrast to another regularly used method, namely method 2 of VanRaden (2008), also described by Yang (2010). This method yields a valid relationship matrix only when the average effect at a locus is proportional to the reciprocal of the square root of expected heterozygosity at that locus (Appendix, Equation A8). So, this method assumes that marker effects are determined by their allele frequency, with larger effects for rarer alleles. For a trait determined by relatively few genes and undergoing directional selection, this assumption may be plausible, since selection acts stronger on causal loci with a larger effect (Haldane 1924; Wright 1931, 1937). It is, however, a very strong assumption in general. Many traits may experience only weak selection, and/or are determined by many genes. In those cases, allele frequency distribution is determined mainly by the interplay of mutation and drift, and a direct relationship between effect size and allele frequency is not expected. Therefore, the assumption of independence between allele frequency and allele substitution effects seems more realistic for most traits. Moreover, when allele substitution effects would depend on allele frequency, effects for exactly the same trait would differ between populations when allele frequencies differ. This makes the interpretation of a genetic correlation estimated using method 2 of VanRaden (2008) rather difficult. Therefore, we advise to use **G** matrices based on method 1 instead of method 2 of VanRaden (2008), especially when multiple populations are considered.

**Table 1.**
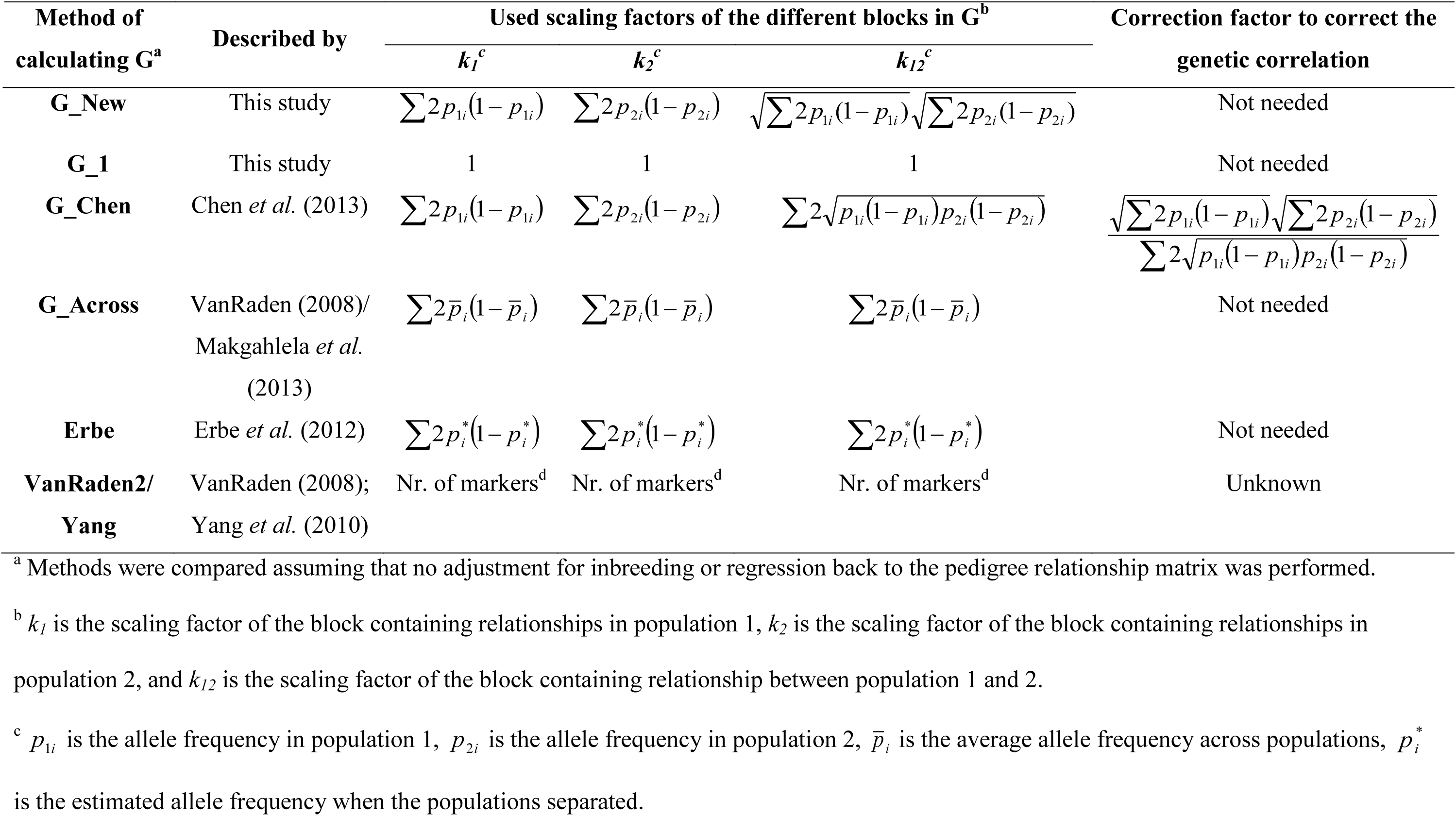
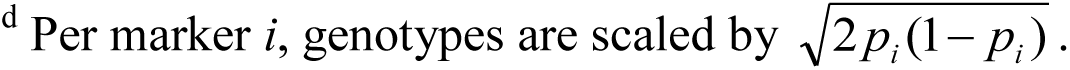
Overview of frequently used method to calculate **G** across populations with scaling and correction factors.

| Method of calculating **G**<sup>a</sup> | Described by | Used scaling factors of the different blocks in **G**<sup>b</sup> |  |  | Correction factor to correct the genetic correlation |
| --- | --- | --- | --- | --- | --- |
| | | $k_1^c$ | $k_2^c$ | $k_{12}^c$ | |
| **G_New** | This study | $\sum 2p_{1i}(1-p_{1i})$ | $\sum 2p_{2i}(1-p_{2i})$ | $\sqrt{\sum 2p_{1i}(1-p_{1i})}\sqrt{\sum 2p_{2i}(1-p_{2i})}$ | Not needed |
| **G_1** | This study | 1 | 1 | 1 | Not needed |
| **G_Chen** | Chen *et al.* (2013) | $\sum 2p_{1i}(1-p_{1i})$ | $\sum 2p_{2i}(1-p_{2i})$ | $\sum 2\sqrt{p_{1i}(1-p_{1i})p_{2i}(1-p_{2i})}$ | $\frac{\sqrt{\sum 2p_{1i}(1-p_{1i})}\sqrt{\sum 2p_{2i}(1-p_{2i})}}{\sum 2\sqrt{p_{1i}(1-p_{1i})p_{2i}(1-p_{2i})}}$ |
| **G_Across** | VanRaden (2008)/<br>Makgahlela *et al.* (2013) | $\sum 2\bar{p}_i(1-\bar{p}_i)$ | $\sum 2\bar{p}_i(1-\bar{p}_i)$ | $\sum 2\bar{p}_i(1-\bar{p}_i)$ | Not needed |
| **Erbe** | Erbe *et al.* (2012) | $\sum 2p_i^*(1-p_i^*)$ | $\sum 2p_i^*(1-p_i^*)$ | $\sum 2p_i^*(1-p_i^*)$ | Not needed |
| **VanRaden2/**<br>**Yang** | VanRaden (2008);<br>Yang *et al.* (2010) | Nr. of markers<sup>d</sup> | Nr. of markers<sup>d</sup> | Nr. of markers<sup>d</sup> | Unknown |

<sup>a</sup> Methods were compared assuming that no adjustment for inbreeding or regression back to the pedigree relationship matrix was performed.
<sup>b</sup> $k_1$ is the scaling factor of the block containing relationships in population 1, $k_2$ is the scaling factor of the block containing relationships in population 2, and $k_{12}$ is the scaling factor of the block containing relationship between population 1 and 2.
<sup>c</sup> $p_{1i}$ is the allele frequency in population 1, $p_{2i}$ is the allele frequency in population 2, $\bar{p}_i$ is the average allele frequency across populations, $p_i^*$ is the estimated allele frequency when the populations separated.
<sup>d</sup> Per marker $i$ , genotypes are scaled by $\sqrt{2p_i(1-p_i)}$ .

In this paper, we assumed that causal loci were known and were used to calculate **G**. In this way, differences in linkage disequilibrium (LD) between markers and causal loci across populations did not affect the results and all genetic variance was explained by **G**. When genomic markers are used to calculate **G,** differences in LD can affect the results, since the LD pattern is known to differ across populations in humans (Sawyer *et al*. 2005) as well as in livestock (e.g., Heifetz *et al*. 2005; Gautier *et al*. 2007; Veroneze *et al*. 2013). This difference in LD is likely to affect the estimated genetic correlation, since it reduces the correlation of marker effects (Gianola *et al*. 2015). Moreover, markers might not explain all genetic variance when there is no complete LD between a causal locus and at least one marker (e.g., Yang *et al*. 2010; Daetwyler *et al*. 2013). This can affect the estimated genetic correlation when the variance explained by the markers shows either a higher or lower genetic correlation than the part not explained (Bulik-Sullivan *et al*. 2015). Therefore, it is difficult to predict the effect of not explaining all genetic variance by markers on the estimated genetic correlation. In a follow-up study, we will investigate the effect of using marker genotypes on the estimated genetic correlation between populations.

### Other approaches to estimate the genetic correlation between populations

We focused on using genomic relationships in a multi-trait model to estimate genetic correlations between populations. Genetic correlations can also be estimated using summary statistics of genome-wide association studies (GWAS; Bulik-Sullivan *et al*. 2015; Brown *et al*. 2016) or using random regression on genotypes (Sørensen *et al*. 2012; Krag *et al*. 2013). The method based on summary statistics of GWAS combines information from different studies and weights estimated marker effects by LD overlap and corresponding *z* score (Bulik-Sullivan *et al*. 2015; Brown *et al*. 2016). This method is beneficial when the costs of collecting enough data are high and data sharing is not possible. It is, however, not known whether this method estimates the correct genetic correlation. The method using random regression on genotypes is equivalent to the multi-trait GREML method used in this study, since both estimate the same additive genetic values when the genotypes are centered and scaled in the same way (Habier *et al*. 2007; VanRaden 2008; Goddard 2009). Variance components estimated with random regression on marker genotypes represent variances of marker effects (Meuwissen *et al*. 2001), similar to **G**_1, when the same centered genotypes are used as input. Hence, random regression on centered genotypes can also be used to estimate genetic correlations between populations. When genotypes for the random regression are centered and scaled, the estimated genetic correlation becomes equal to the estimated genetic correlation using **G** based on method 2 of VanRaden (VanRaden 2008; Yang *et al*. 2010). Therefore, the interpretation of this estimated genetic correlation remains unclear as well.

### Importance of the genetic correlation between populations

The genetic correlation between populations is an important parameter for genomic prediction, since it determines the usefulness of combining information from multiple populations. A low genetic correlation means that it is very unlikely that combining populations will increase the accuracy of estimated genetic values. Therefore, the genetic correlation partly determines the accuracy of across- or multi-population genomic prediction. For predicting the accuracy in those scenarios, an accurate estimation of genetic correlations is essential (Wientjes *et al*. 2015; Wientjes *et al*. 2016). For predicting response to selection, both the accuracy as well as current genetic variances are needed (Falconer and Mackay 1996). Even though the accuracy of estimated genetic values is quite consistent across methods for calculating **G** (Makgahlela *et al*. 2013, 2014; Lourenco *et al*. 2016), for estimating genetic (co)variances and correlations it is important to use the **G**_New matrix.

### Genetic correlation versus genic correlation

The genetic correlation is defined based on additive genetic (co)variances. Under selection, however, additive genetic (co)variances change over generations, since selection creates transient gametic phase disequilibrium (i.e., correlations between allele substitution effects at different loci). This process is also known as the Bulmer effect (Bulmer 1971). Therefore, genetic (co)variances and correlations depend not only on the genetic background of the traits, but also on transient processes like the type and intensity of selection. Apart from additive genetic (co)variances, quantitative genetics also describes genic (co)variances (e.g., Bulmer 1980; Bulmer 1989), defined as the additive genetic (co)variance in the absence of gametic phase disequilibrium. In contrast to genetic variances, genic variances are independent from selection and are always equal to twice the Mendelian sampling variance (Hill 2014). In analogy to genic (co)variances, genic correlations can be defined as well. We believe that genic correlations are more relevant than additive genetic correlations, since genic correlations are not influenced by transient processes and, therefore, more constant across generations.

In our simulation study, allele substitution effects were randomly sampled, so no transient gametic phase disequilibrium was present and genic (co)variances were equal to the additive genetic (co)variances. In all situations, genic variances can be estimated when the base population of the relationship matrix is unselected and phenotypic records on which selection decisions are based are available (Henderson 1985). It is also shown that even when phenotypic records from the base population are absent, the genic variance can be estimated when phenotypic records for several generations are available and the base population is unselected (Henderson 1985; Van der Werf and de Boer 1990). It can be expected that as long as several generations of phenotypic data is available in combination with the relationships between all those individuals, variances are corrected for selection and effectively genic variances are estimated. Therefore, genic correlations can likely be calculated using **G**_New, provided that data is available for several generations.

## Conclusion

The properties of the genomic relationship matrix affect estimates of genetic variances within as well as genetic correlations between populations. For estimating current genetic variances, allele frequencies of the current population should be used to calculate relationships within that population. For estimating genetic correlations between populations, scaling factors of the different blocks of the relationship matrix, based on method 1 of VanRaden (2008), should fulfill the property 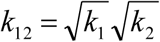. When this property is not fulfilled, the estimated genetic correlation can be corrected by multiplying the estimate by 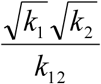. In this study we present a genomic relationship matrix, **G**_New, which directly ^k^12 results in current genetic variances as well as genetic correlations between populations.

## ACKNOWLEDGMENTS

This research is supported by the Netherlands Organisation of Scientic Research (NWO) and the Breed4Food consortium partners Cobb Europe, CRV, Hendrix Genetics and TopigsNorsvin.

## APPENDIX

The **G** matrix based on method 2 of VanRaden (2008) and Yang *et al*. (2010), **G_**VR2, weights markers by the reciprocal of the square root of the variance of its genotypes. In this Appendix, it is shown that this is only correct under the assumption that the variance of the average effect (α) at a locus, say l, is inversely proportional to expected heterozygosity at that locus,

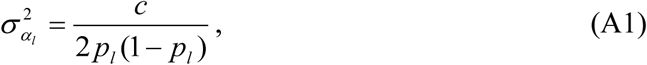
where *c* is a constant, and *p_l_* the allele frequency at locus *l*.

Consider the single-trait mixed model **y = Xb + Za + e**, where **a** is the vector of random additive genetic effects, with 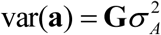. This mixed model is valid only when 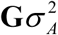 indeed represents the covariances between additive genetic effects (*A*) of individuals. This requires that

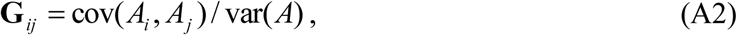
where *i* and *j* are individuals.

By definition, the additive genetic effect of an individual is the sum of the average effects at its loci, weighted by the centred allele count (Fisher 1918; Falconer and Mackay 1996),

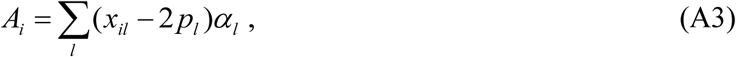
where *x_ij_* is the allele count of individual *i* at locus *l*, taking values 0, 1 or 2. Thus

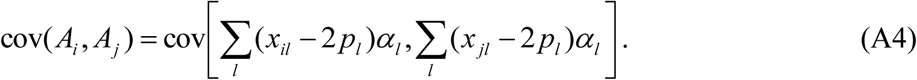

For the genic covariance, the (*x_il_* –2*p_l_*) *α_l_* terms are independent between loci by definition (Bulmer 1971), so that the covariance reduces to

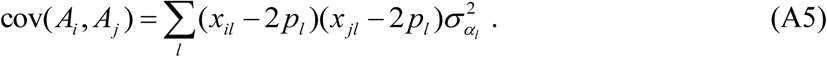

Substituting the relationship between average effects and allele frequency given by Equation A1 yields

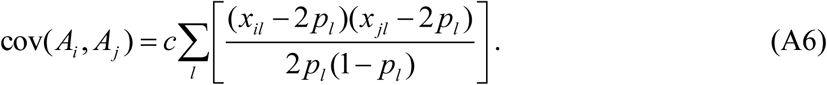

Analogously, the genic variance equals

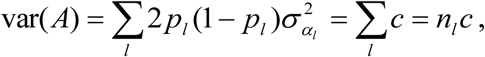
where *n_l_* is the number of loci. Finally, from Equation A2,

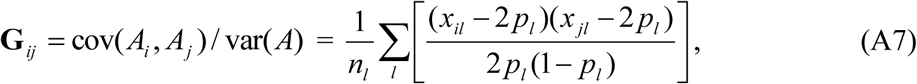
which is **G**_VR2. Thus obtaining **G**_VR2 requires Equation A1.

Hence, **G**_VR2 is valid under the assumption that the magnitude of the average effect at a locus is proportional to the reciprocal of the square root of expected heterozygosity at that locus,

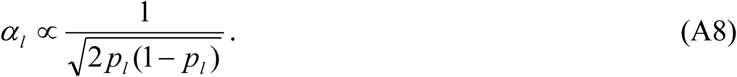

Equation A7 shows that elements of **G**_VR2 are the genome-wide average of the correlations at individual loci; the term in square-brackets is the correlation between additive genetic effects at locus *l*, and the sum of these terms is divided by the number of loci. Thus **G**_VR2 may have been motivated as the genome-wide average of relationships at individual loci.

However, relatedness refers to the correlation between the total additive genetic effects of individuals (Equation A2), which are sums of additive genetic effects at individual loci. In general, the correlation between sums does not equal the average correlation between components of the sums,

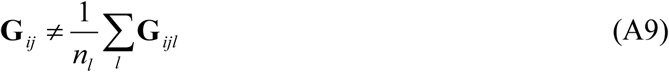
but is defined as the ratio of the covariance and variance of the sum,

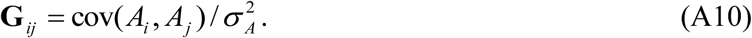
Equations A9 and A10 are only equal to each other under the assumption given in Equation A1.

## LITERATURE CITED

Bohren, B. B., W. G. Hill and A. Robertson, 1966 Some observations on asymmetrical correlated responses to selection. Genet. Res. 7: 44–57.

Brown, B. C., C. J. Ye, A. L. Price and N. Zaitlen, 2016 Transethnic genetic-correlation estimates from summary statistics. Am. J. Hum. Genet. 99: 76–88.

Bulik-Sullivan, B., H. K. Finucane, V. Anttila, A. Gusev, F. R. Day, et al., 2015 An atlas of genetic correlations across human diseases and traits. Nat. Genet. 47: 1236–1241.

Bulmer, M., 1989 Maintenance of genetic variability by mutation-selection balance: a child’s guide through the jungle. Genome 31: 761–767.

Bulmer, M. G., 1971 The effect of selection on genetic variability. Am. Nat. 105: 201–211.

Bulmer, M. G., 1980 The mathematical theory of quantitative genetics. Clarendon Press., Oxford.

Chen, L., F. Schenkel, M. Vinsky, D. Crews and C. Li, 2013 Accuracy of predicting genomic breeding values for residual feed intake in Angus and Charolais beef cattle. J. Anim. Sci. 91: 4669–4678.

Daetwyler, H. D., M. P. L. Calus, R. Pong-Wong, G. De los Campos and J. M. Hickey, 2013 Genomic prediction in animals and plants: Simulation of data, validation, reporting, and benchmarking. Genetics 193: 347–365.

De Candia, T. R., S. H. Lee, J. Yang, B. L. Browning, P. V. Gejman, et al., 2013 Additive genetic variation in schizophrenia risk is shared by populations of African and European descent. Am. J. Hum. Genet. 93: 463–470.

Erbe, M., B. J. Hayes, L. K. Matukumalli, S. Goswami, P. J. Bowman, et al., 2012 Improving accuracy of genomic predictions within and between dairy cattle breeds with imputed high-density single nucleotide polymorphism panels. J. Dairy Sci. 95: 4114–4129.

Falconer, D. S., 1952 The problem of environment and selection. Amer. Nat. 86: 293–298.

Falconer, D. S. and T. F. C. Mackay, 1996 Introduction to quantitative genetics. Pearson Education Limited, Harlow.

Fisher, R. A., 1918 The Correlation between relatives on the supposition of Mendelian inheritance. Trans. Roy. Soc. Edinburgh 52: 399–433.

Fisher, R. A., 1930 The genetical theory of natural selection: a complete variorum edition. Oxford University Press.

Gautier, M., T. Faraut, K. Moazami-Goudarzi, V. Navratil, M. Foglio, et al., 2007 Genetic and haplotypic structure in 14 European and African cattle breeds. Genetics 177: 1059–1070.

Gianola, D., G. De los Campos, M. A. Toro, H. Naya, C.-C. Schön, et al., 2015 Do molecular markers inform about pleiotropy? Genetics 201: 23–29.

Goddard, M. E., 2009 Genomic selection: Prediction of accuracy and maximisation of long term response. Genetica 136: 245–257.

Habier, D., R. L. Fernando and J. C. M. Dekkers, 2007 The impact of genetic relationship information on genome-assisted breeding values. Genetics 177: 2389–2397.

Haldane, J. B. S., 1924 A mathematical theory of natural and artificial selection—I. Trans. Camb. Phil. Soc. 23: 19–41.

Harris, B. L. and D. L. Johnson, 2010 Genomic predictions for New Zealand dairy bulls and integration with national genetic evaluation. J. Dairy Sci. 93: 1243–1252.

Hayes, B. J., P. M. Visscher and M. E. Goddard, 2009 Increased accuracy of artificial selection by using the realized relationship matrix. Genet. Res. 91: 47–60.

Heifetz, E. M., J. E. Fulton, N. O’Sullivan, H. Zhao, J. C. M. Dekkers, et al., 2005 Extent and consistency across generations of linkage disequilibrium in commercial layer chicken breeding populations. Genetics 171: 1173–1181.

Henderson, C. R., 1985 Best linear unbiased prediction using relationship matrices derived from selected base populations. J. Dairy Sci. 68: 443–448.

Hill, W. G., 2014 Applications of population genetics to animal breeding, from Wright, Fisher and Lush to genomic prediction. Genetics 196: 1–16.

Karoui, S., M. Carabaño, C. Díaz and A. Legarra, 2012 Joint genomic evaluation of French dairy cattle breeds using multiple-trait models. Genet. Sel. Evol. 44: 39.

Krag, K., N. A. Poulsen, M. K. Larsen, L. B. Larsen, L. L. Janss, et al., 2013 Genetic parameters for milk fatty acids in Danish Holstein cattle based on SNP markers using a Bayesian approach. BMC Genet. 14: 79.

Legarra, A., 2016 Comparing estimates of genetic variance across different relationship models. Theor. Popul. Biol. 107: 26–30.

Lehermeier, C., C.-C. Schön and G. De los Campos, 2015 Assessment of genetic heterogeneity in structured plant populations using multivariate whole-genome regression models. Genetics 201: 323–337.

Lourenco, D. A. L., S. Tsuruta, B. O. Fragomeni, C. Y. Chen, W. O. Herring, et al., 2016 Crossbreed evaluations in single-step genomic best linear unbiased predictor using adjusted realized relationship matrices. J. Anim. Sci. 94: 909–919.

Makgahlela, M. L., I. Strandén, U. S. Nielsen, M. J. Sillanpää and E. A. Mäntysaari, 2013 The estimation of genomic relationships using breedwise allele frequencies among animals in multibreed populations. J. Dairy Sci. 96: 5364–5375.

Makgahlela, M. L., I. Strandén, U. S. Nielsen, M. J. Sillanpää and E. A. Mäntysaari, 2014 Using the unified relationship matrix adjusted by breed-wise allele frequencies in genomic evaluation of a multibreed population. J. Dairy Sci. 97: 1117–1127.

Meuwissen, T. H. E., B. J. Hayes and M. E. Goddard, 2001 Prediction of total genetic value using genome-wide dense marker maps. Genetics 157: 1819–1829.

Olson, K. M., P. M. VanRaden and M. E. Tooker, 2012 Multibreed genomic evaluations using purebred Holsteins, Jerseys, and Brown Swiss. J. Dairy Sci. 95: 5378–5383.

Powell, J. E., P. M. Visscher and M. E. Goddard, 2010 Reconciling the analysis of IBD and IBS in complex trait studies. Nat. Rev. Gen. 11: 800–805.

Sawyer, S. L., N. Mukherjee, A. J. Pakstis, L. Feuk, J. R. Kidd, et al.,2005 Linkage disequilibrium patterns vary substantially among populations. Europ. J. Hum. Genet. 13: 677–686.

Sørensen, L. P., L. Janss, P. Madsen, T. Mark and M. S. Lund, 2012 Estimation of (co)variances for genomic regions of flexible sizes: application to complex infectious udder diseases in dairy cattle. Genet. Sel. Evol. 44: 18.

Speed, D. and D. J. Balding, 2015 Relatedness in the post-genomic era: is it still useful? Nat. Rev. Genet. 16: 33–44.

Strandén, I. and O. F. Christensen, 2011 Allele coding in genomic evaluation. Genet. Sel. Evol. 43: 25.

Van der Werf, J. and I. de Boer, 1990 Estimation of additive genetic variance when base populations are selected. J. Anim. Sci. 68: 3124–3132.

VanRaden, P. M., 2008 Efficient methods to compute genomic predictions. J. Dairy Sci. 91: 4414–4423.

Veroneze, R., P. S. Lopes, S. E. F. Guimarães, F. F. Silva, M. S. Lopes, et al., 2013 Linkage disequilibrium and haplotype block structure in six commercial pig lines. J. Anim. Sci. 91: 3493–3501.

Visscher, P. M., G. Hemani, A. A. E. Vinkhuyzen, G.-B. Chen, S. H. Lee, et al., 2014 Statistical power to detect genetic (co)variance of complex traits using SNP data in unrelated samples. PLoS Genet 10: e1004269.

Wientjes, Y. C. J., R. F. Veerkamp, P. Bijma, H. Bovenhuis, C. Schrooten, et al., 2015 Empirical and deterministic accuracies of across-population genomic prediction. Genet. Sel. Evol. 47: 5.

Wientjes, Y. C. J., P. Bijma, R. F. Veerkamp and M. P. L. Calus, 2016 An equation to predict the accuracy of genomic values by combining data from multiple traits, breeds, lines, or environments. Genetics 202: 799–823.

Wright, S., 1931 Evolution in Mendelian populations. Genetics 16: 97–159.

Wright, S., 1937 The distribution of gene frequencies in populations. Proc. Nat. Acad. Sci. USA 23: 307–320.

Yang, J., B. Benyamin, B. P. McEvoy, S. Gordon, A. K. Henders, et al., 2010 Common SNPs explain a large proportion of the heritability for human height. Nat. Genet. 42: 565–569.

Yang, J., A. Bakshi, Z. Zhu, G. Hemani, A. A. E. Vinkhuyzen, et al., 2015 Genetic variance estimation with imputed variants finds negligible missing heritability for human height and body mass index. Nat. Genet. 47: 1114–1120.

